# Protein Phosphatase 2A Orchestrates Mitochondrial Dynamics via MAPK Signaling in heart

**DOI:** 10.64898/2026.02.26.708402

**Authors:** Dachuan Dong, Yin Zhang, Liangyuan Li, Hongwei Fan, Tao Jin, Xiang Gao, Zhao Zhang

**Author notes:** Division of Endocrinology, Gerontology, and Metabolism, School of Medicine, Stanford University, Stanford 94305, United States (Dachuan Dong). State Key Laboratory of Medical Neurobiology and MOE Frontiers Center for Brain Science, School of Basic Medical Science, Fudan University, Shanghai 200032. China. These authors contributed equally to this study (Dachuan Dong, Yin Zhang).

## Abstract

Heart as one high ATP consuming organ accounts for 5% of the total oxygen demands. The central question of heart health is how mitochondria fit its needs. Impaired mitochondrial dynamics (fission and fusion) have been observed in failing heart, but whether and how phosphorylation events involved in mitochondrial quality control are still imperceptive. The phosphatase 2A catalytic subunit α (PP2A cα) cardiac-specific knockout mouse (KO), which exhibited a hypertrophic cardiomyopathy phenotype, was studied. We profiled the pattern of morphological and functional alteration of cardiac mitochondria that appeared during postnatal development. Increased heterogeneity of mitochondria and a decreased ATP yield was displayed. Notably, a fission procedure escalated. To illustrate the protagonist of the mitochondrial dynamics, we applied a high-throughput spectrometry-based phosphoproteomic screening following by GO and KEGG pathway annotations for 788 phosphosites, accounting for 90 proteins. Results suggested that the MAPK signaling may be a predominant factor associated with those mitochondrial alternations in KO hearts. Furthermore, we identified hyperphosphorylated ERK2 accumulated into the nucleus regarding PP2Acα depletion. Consequently, *Fis1* expression was accelerated at the transcriptional level which facilitated recruitment of Drp1 onto the outer mitochondrial membrane. The mitochondrial fission towards shifting led to excessed mitophagy and is considered the culprit in early mortality. These findings are indicative of the fundamental role of PP2A in mitochondrial dynamics regulation and cardiomyopathy progression. During the progression of heart failure, the phospho-regulation of ERK2 could be a novel therapeutic approach to prevent or attenuate adverse hypertrophic cardiomyopathy.

## 1 Introduction

The mammalian heart is a low energy storage but a mitochondria-enriched organ. One-third of the myocardial mass is constituted by mitochondria that are capable of producing a high amount of available energy to sustain normal contractile function[1]. Under physiological conditions, the heart is fully aerobic, while over 90% of the ATP production is via mitochondrial oxidative phosphorylation[2, 3], At birth, in response to a switch from placental to the pulmonary circulation, the heart commences its aerobic metabolism that is accompanied by a complex remodeling process from fetal to an adult state. In myocardial cells, this transition requires not only a metabolic switch to initiate aerobic metabolism[4, 5], but also a continuous maturation of mitochondria by dramatically increasing their size and function, together with rearrangement of the organelles throughout the myocyte[6, 7]. Emerging evidence reveals that postnatal mitochondrial maturation in the mammalian heart involves a profound remodeling event termed as mitochondrial dynamics by undergoing coordinated fusion and fission[8]. Of note, mitochondrial dynamics is not just an indication of the change in mitochondrial morphology. It also functions as a tight link between the maturation of intracellular energy pathways and cell architecture to ensure proper postnatal heart transition[9]. Moreover, the dynamic alteration of mitochondrial morphology through fusion and fission events allows the exchange of mitochondrial content and segregation of terminally damaged mitochondria. This enables degradation by selective autophagy called mitophagy. As such, the homeostasis of fusion and fission are established as the cornerstone of mitochondrial quality control, while abnormal mitochondrial dynamics has implicated in heart diseases[10–12]. Although mito-dynamics has attracted increased attention in recent years, its potential role in this process during postnatal maturation of the heart has not been completely characterized.

Reversible phosphorylation regulated by kinases and protein phosphatases (PPs) is one of the most prevalent post-translational modifications. In the heart, more than 90% of dephosphorylation events are conducted by PP1 and PP2A. The latter is ubiquitously expressed in tissues with one of the most abundant Ser/Thr phosphatases in nature. The holoenzyme of PP2A is a heterotrimer composed by scaffolding A subunit, alternative regulatory B subunit, and a catalytic C subunit[13]. In mammalian cardiomyocytes, two isoforms of PP2Ac (cα and cβ) have been identified and PP2Acα is the predominant isoform[14]. To address the function of PP2A in heart tissues, several genetic and functional studies have been established[15–17]. Our previous work showed that conditional inactivation of cardiac PP2Acα in mice led to shortening lifespan accompanied with impaired cardiac function[18]. Consistent with our observations, several murine models with mitochondrial impairments exhibit early mortalities during postnatal development as well[16, 19]. Based on this evidence, we hypothesized that PP2Acα might be engaged in mitochondrial dynamics, contributing to heart dysfunction at early postnatal development.

In this study, we used mice with cardiac-specific deletion of PP2Acα intended to provide a fundamental understanding of how phosphorylation events orchestrate mitochondrial dynamics and heart development. Our findings revealed that the homeostasis of cardiac phosphorylation plays essential roles in mitochondrial quality control during the early stage of postnatal development. In addition, our observations suggested that suppressing mitochondrial fission driven by MAPK or inhibitors targeting this process might be a promising therapeutic strategy for heart failure.

## 2 Results

### 2.1 Conditional depletion of PP2Acα impaired mitochondrial architecture and oxidative performance

The cardiac-specific *Ppp2ca*^-/-^ knockout mouse (KO) exhibited hypertrophic cardiomyopathy during postnatal development since Cre recombinase expressed since P6.5. As the consequence of severely impaired ejection function, the KO mice suddenly died around P12 (Fig. S.1).

To answer the question of whether the mitochondrial impairment is involved in a sudden death in KO mice, we investigated the ultrastructure of the heart by using transmission electron microscopy (TEM). In Ctrl mouse hearts, the counts of mitochondria increased gradually during postnatal development, while the mitochondrial size and aspect ratio (AR, length of long axis/short axis) had no significant changes. The mitochondria in the KO hearts at P7 exhibited rod-shaped structure (Fig. 1A, bottom left), reflected by significant increases of aspect ratio (4.6 vs 1.7,). In addition, most longitudinal mitochondria turned to round particles (AR value 1.1) and the relative contents of mitochondria increased (1.5-fold change) but their size significantly reduced (0.4μm^2^ vs 1.0μm^2^), implying there were profound changes in mitochondrial dynamics from P7 to P9 in KO cardiomyocytes. These spherical intermyofibrillar mitochondria still maintained a relatively integrated architecture at P9. Notably, mitochondria in KO hearts at P11 swelled extremely, some of which developed obvious broken-holes on the membranes. The cristaes collapsed into fragments, and the giant burbling cavity occurred in the mitochondrial lumen (Fig. 1A, bottom right). With aggravation of the damage, the arrangement of mitochondria in KO cardiomyocytes was no longer in a regular pattern (Fig. 1A and 1B).

**FIG. 1.**
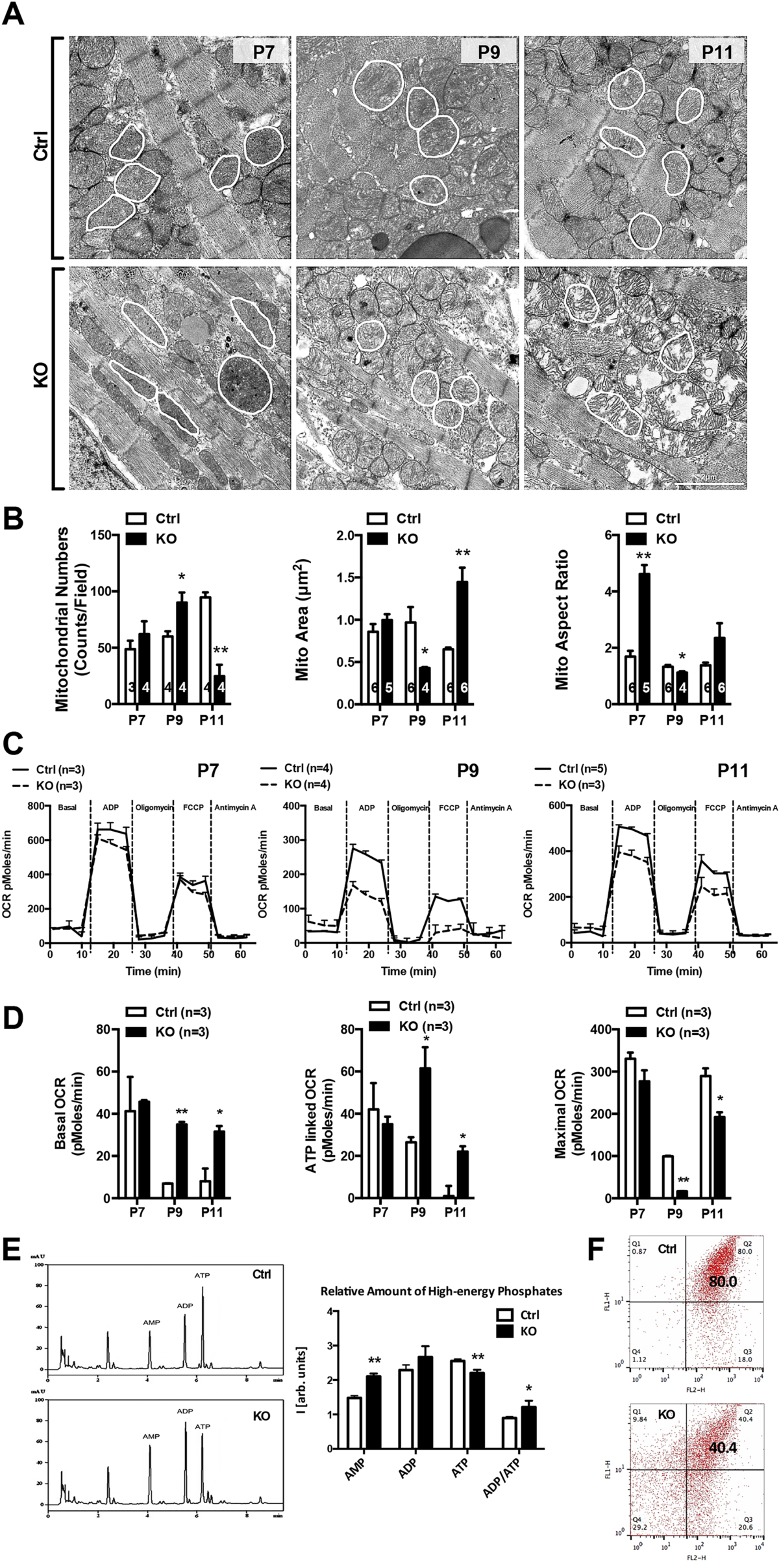
Progressive cardiac mitochondrial structural and functional impairments developed in postnatal PP2Acα ablation mouse hearts. (A), PP2Acα deficiency evoked mitochondria dysmorphometry in mouse hearts. Representative transmission electron microscopy images of left ventricular on P7 to P11. (B), Quantitative mitochondrial numbers (counts to per field, left), mitochondrial aspect ratio (middle) and mitochondrial area (right). (C), Representative graph showing the mitochondrial oxygen consumption rates (OCRs) of cardiomyocytes, from P7-P11 mouse hearts, were tested by Seahorse XF24 analyzer. Addition of ADP, oligomycin, FCCP and antimycin A is indicated. (D), Quantification of basal OCR, ATP linked OCR and Maximal OCR data from (C). (E), Original traces from HPLC experiments with standards solution containing AMP, ADP, and ATP are shown. Signal intensity was representative in arbitrary units of absorption at 260nm (left). Quantitative concentrations of high-energy phosphates (right). (F), The mitochondrial membrane potential was measured using flow cytometry. Isolated mitochondrial monitored by JC-1. All data reported as mean±S.E., \**P*<0.05, \*\**P*<0.01 vs Ctrl group. n= indicated numbers of mice.

To evaluate the impact of mitochondrial dynamic changes on respiratory capacity, we assessed the oxygen consumption rate (OCR). Results showed that mitochondrial functions were stable in P7 myocytes for either group, while at a final stage of the PP2Acα ablation, the mitochondria in KO cardiomyocytes showed an enhanced basal OCR level and a lowered maximum respiratory rate (Fig. 1C and 1D). These data suggested that lacking cardiac PP2A is sufficient to compromise the mitochondrial oxidative functions by fetal genetic reprogramming followed by increased cell size. Next, we examined the effects of PP2Acα deficiency on ATP generation by high-performance liquid chromatography (HPLC) assay. Indeed, KO hearts had significantly less high-energy phosphates than the Ctrl and this paralleled a remarkable decline in conversion capability from ADP to ATP (ADP/ATP ratio was 0.89 vs. 1.21 in Ctrl and KO hearts, respectively; Fig. 1E), demonstrating a remarkably decreased ATP yields in KO hearts (86.17% of the Ctrl). In addition to the functional impairment of mitochondria, a significant disruption of mitochondrial membrane potential (MMP, Δψm) was evident in KO mice in comparison with the Ctrl (Fig. 1F).

### 2.2 A phosphoproteomic perspective of putative interaction networks for PP2A

PP2A is recognized as a dynamic and versatile protein phosphatase which is involved in many signaling pathways and cellular functions. Therefore, the impact of PP2Acα deficiency might cause profound multi-level changes. Here, we focused on the post-translational level. In order to identify which major phosphorylated molecule contributes to myocardial metabolism, we performed a phosphoproteomic screening looking for the key element that orchestrates mitochondrial dynamics in developmental mouse hearts.

Heart tissues were freshly collected and rigidly homogenized before subjecting to an LCMS-IT-TOF based Perfinity iDP proteomics, where enrichment of phosphorylated peptides with iTRAQ labeling was used to calculate the differences of phosphorylation status between Ctrl and KO hearts. A total of 507 unique phosphopeptides corresponding to 788 valid phosphorylation sites on 90 distinct proteins (supplementary material) were identified with a confidence interval for the identification set at 95%, among which 303 peptides on 464 sites were enhanced and 204 peptides on 324 sites were decreased. Considering the PP2A as one protein phosphatase, we assume that the most likely outcome of PP2Acα deficiency should be higher phosphorylation levels while the events of the decreased level of phosphorylation might not be affected directly by PP2A. Therefore, we sieved putative candidates narrowing down to a 2-fold up-regulated level (Log_2_FC≥1), and then 37 phosphorylation sites on 15 non-redundant proteins that met our criteria stood out for further analysis (Fig. 2A). Results of protein-protein interaction (PPI) mapping indicated that PP2Acα substantially connected with CDC25, CHK2, MAPK1, PP1B, IKK, and others. Following by Gene Ontology (GO) functional annotations, PPI datasets were involved in a wide variety of biological processes that were roughly categorized into six portions (top panel in Fig. S.2), such as metabolic pathway, DNA damage and repair, development process, protein transport, signal transduction, and transcriptional regulation. Among those metabolic changes, accounting for 46.3% in total biological events, became the key outcomes in response to PP2Acα deficiency (right bottom in Fig. S.2). Furthermore, these proteins mainly distribute in the cytoplasm and function as the regulators for phosphorylation activity (bottom-left in Fig. S.2)

**FIG. 2.**
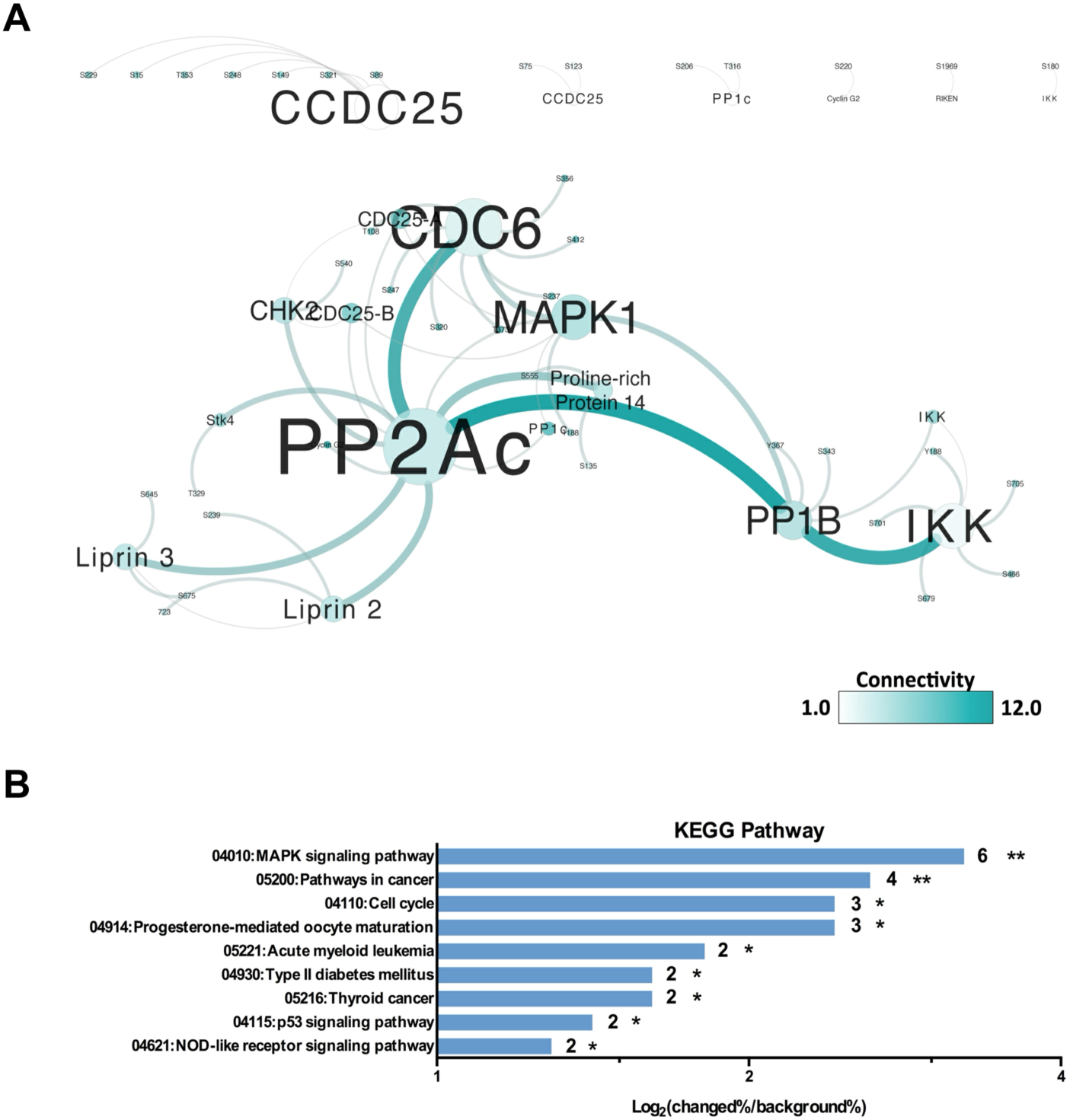
Identifying phosphorylated proteins influenced by PP2Acα. (A), Schematic for phosphoproteomics-based identification sets. (B), Identification of regulated phosphorylation in response to PP2Ac depletion. The geometric mean of iTRAQ spectral peak height ratios for replicate samples was plotted as a function of the sum of the geometric means of all iTRAQ spectral peak heights. The 95% acceptance region was calculated (grey curve). (C), Interactions between hyper-phosphorylated proteins identified by phosphoproteomics. The connectivity was performed by size and coloration. (D), An overview of signaling pathways regulated by PP2Ac. PANTHER Pathway terms were extracted for each phosphor-peptides that changed significantly with PP2Acα.

Next, we used the Panther database KEGG pathway annotations[20, 21] to analyze the impact on the underlying pathways in response to PP2Acα ablation. A total of 9 signaling pathways were identified, among which a large portion were involved in the MAPK signaling pathway (Fig. 2B). A comprehensive method of analyzing the results of phosphoproteomic screening suggested that PP2Acα deficiency closely contacted with MAPK pathway. Additionally, functional network analysis also highlighted the most likely protein involved in mitochondrial quality control is MAPK1, also known as ERK2.

### 2.3 PP2A directly de-phosphorylates ERK2 at dual phosphor-sites

Considering MAPKs as one of the most prominent pathways impacted by PP2Acα deficiency and EKR2 as the main molecule involved in, we expect the observed putative associations to represent the proteomic perspective, which was necessary for us to gain an understanding of complexes that PP2Acα may associate with. Primarily, to distinguish the interaction affinity, we applied a series of lysis buffers, a range of mild NP-40 buffer (1% NP-40) to stringent RIPA buffer (0.1% SDS, 1% Triton X-100), to treat the supernatant obtained from KO and control hearts. As expected, immunoaffinity precipitation exhibited a tight-association between PP2Acα and ERK2. The association was not disrupted even in the presence of a harsh detergent (Fig. 3A), implying that the PP2A manipulated the phosphorylation status of ERK2 directly without mediators. As previous phosphoproteomic reports have shown, ERK2 contains multiple phosphorylated sites, among which, modifiable residues (T179, T183, Y185, and T188) clustered at the PKinase domain region (23-311aa) responding to its kinase activity (Fig. 3B) demonstrated enhanced phosphorylation on phosphoproteomic screening. The comparison of ERK2 orthologs via NCBI Standard Protein BLAST showed that the modification residues are highly evolutionarily conserved in vertebrates (Fig. 3C). Additionally, using the PONDR (Predictor Of Naturally Disordered Regions) algorithm, we determined these phosphorylated residues of ERK2 are within a topological stable region (Fig. 3D). Percussive research indicated the structural stability and activity of ERK2 are tightly controlled at T183 and Y185[22–24]. Herein, we focused on the dual-phosphorylation sites in the following studies.

**FIG. 3.**
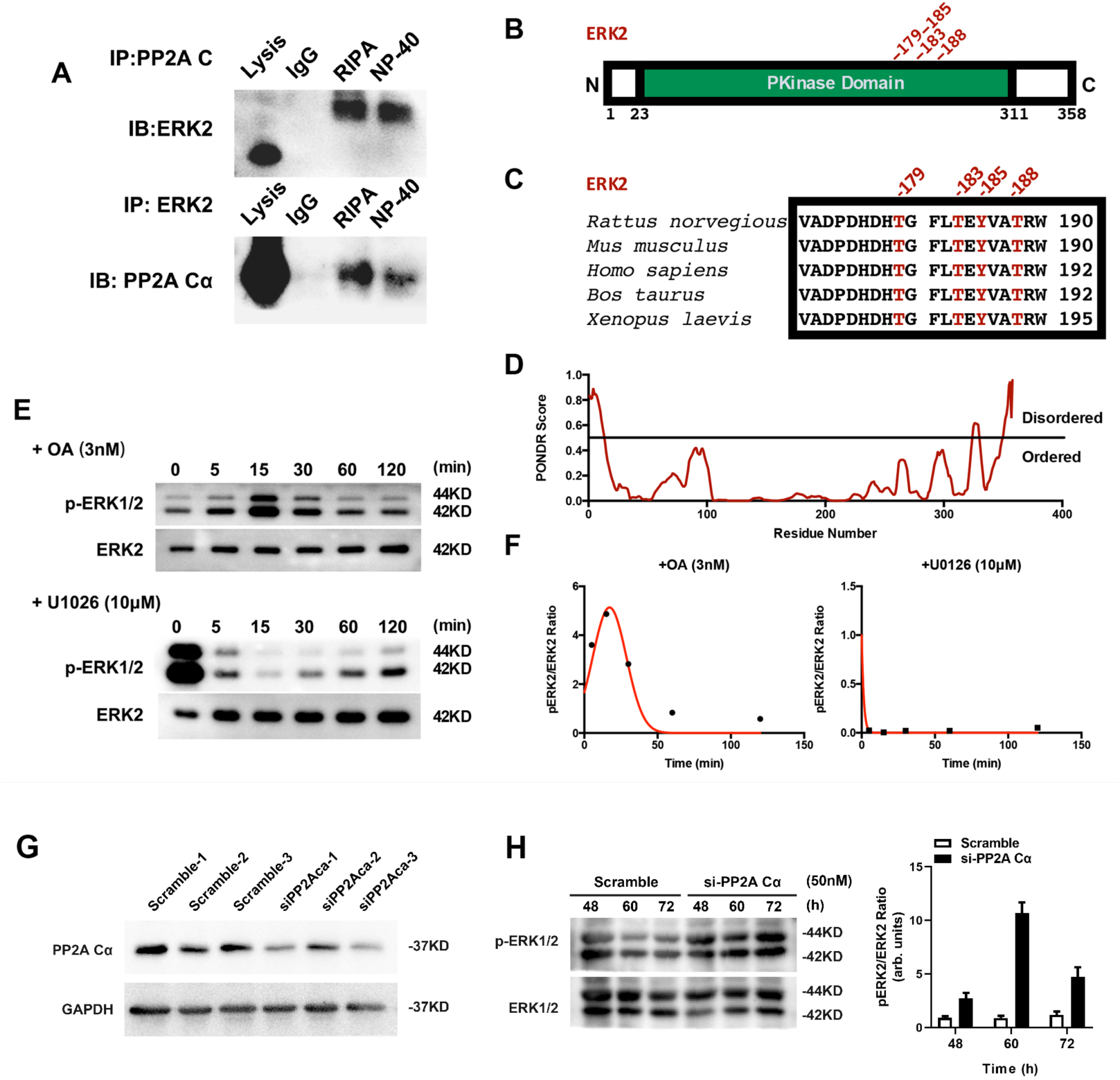
PP2Acα interact with ERK2. (A), Association of PP2Acα with ERK2 in situ. Followed by lysis and immunoprecipitation with anti PP2Acα and anti ERK2 antibodies. (B), ERK2 is phosphorylated at multiple residues within a PKinase domain region: T179, T183, Y185 and T188. (C), Phosphorylated ERK2 residues (red) are evolutionarily conserved among metazoan species. Sequence alignment was generated by NCBI standard protein BLAST. (D), ERK2 is predicted to be natively ordered at its PKinase domain by the PONDR algorithm. (E), Phosphorylation of ERK2 is an early response event in H9c2 cell line, treated by OA or U0126. (F), Quantitated phosphorylation ratio of ERK2 in (E). Red lines represent peak integration with Gaussian fitting. (G), The effectiveness of the si-PP2A Cα at 48hs. (H,) Phosphorylation of ERK2 in H9c2 cells after si-PP2A Cα transfection.

Surprisingly, in P11 hearts, both the expression and modification level of three MAPKs members have no changes except the mildly decreased phosphor-Jnk1/2/3 (Fig. S3A). In regard to this, we detected the changes of ERK2 at different time points and found that the phosphorylation level of p-ERK2^T183/Y185^ in KO hearts was strikingly elevated at P7, and then returned to a relatively steady level at P9 and P11 (Fig. S3B) as compare to Ctrl. To confirm this observation, we treated H9c2 cells with Okadaic Acid (OA, 5nM), a specific PP2A inhibitor[25, 26] and U0126 (10μM), an inhibitor of MEK1/2[27, 28] which catalyzes the activation of the effector MAP kinases ERK1/2, respectively. Consistent with observations *in vivo*, the response of p-ERK2^T183/Y185^ to the treatments occurred rapidly. The phosphorylation level of ERK2 reached its climax at 15 min, and then gradually declined back to basal level after 30 min (Fig. 3E, F). In order to better understand the impact of PP2A on ERK, H9c2 cells were transfected with si-PP2A Cα to mimic the KO condition (Fig. 3G). Similar to OA treatments, the level of p-ERK2^T183/Y185^ went up and reached its vertex at 60h after transfection then gradually fade away (Fig. 3H). Meanwhile, sustained OA treatment (5nM, 24h) resulted in a reduction of PP2A expression (Fig. S3C). Taken together, these results suggested that ablation of PP2Acα predominantly impacted the MAPK signaling pathway, especially the hyperphosphorylation of ERK2, at the early stage of gene ablation.

### 2.4 Subcellular distributions of ERK2 determined by its phosphorylation level

As described before, the PP2A strongly modulated the phosphorylation level of ERK2 through direct interaction. Phosphorylated ERK2 usually clumped and dimerization status determined its subcellular localization. Therefore, we speculated that PP2Acα deletion or inhibition may impact on ERK2 translocation. To explore this possibility, cardiomyocytes were isolated from P7 to P11 mouse hearts and labeled with an anti-p-ERK2^T183/Y185^ antibody. It was intriguing to find that the p-ERK2 ^T183/Y185^ are ubiquitously distributed in the cardiomyocytes during postnatal development. Notably, p-ERK2 ^T183/Y185^ is widespread in the cytoplasm in early stage (P7) control cardiomyocytes. In contrast, in KO cardiomyocytes, high intensity of the fluorescent signal of p-ERK2 ^T183/Y185^ appeared predominately in the nuclei (Fig. 4A and 4B). However, after P7, there are no significant differences in subcellular distribution for both groups (Fig. 4A and 4B). We took a further step to confirm this observation in H9c2 cells in the presence of OA or U0126, respectively, on a time scale. Corresponding results displayed that a peak of p-ERK2^T183/Y185^ nuclear accumulation occurred at 15min in the presence of PP2A and then reduced back to basal level gradually. Treatments with either OA or U0126 effectively abrogated the nuclear transportation of p-ERK2^T183/Y185^, recapturing the data from mouse cardiomyocytes (Fig. 4C). These findings demonstrated that the PP2A was involved in ERK2 subcellular distribution through dual-phosphorylation sites of ERK2 at T183/Y185.

**FIG. 4.**
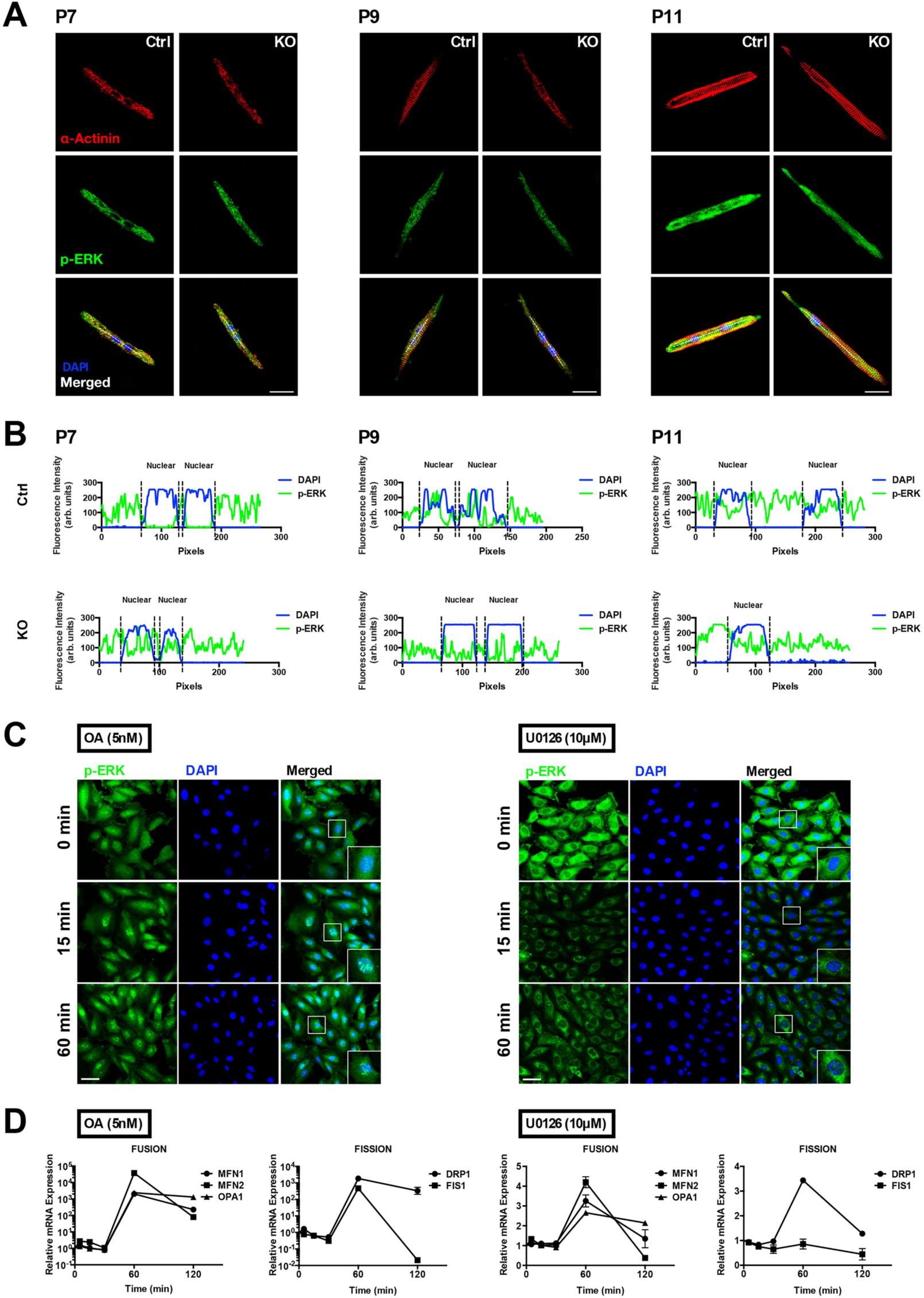
Phosphorylation of ERK2 determined its subcellular localization. (A), Cardiac cells, isolated from KO mice and their paired control on P9-P11, was staining by anti α-Actinin (Red); anti p-ERK2^T183/Y185^(Green) and DAPI (Blue), Scale bar=20µm. (B), Quantitative fluorescence intensity illustrated the merged image in (A). (C), Cultured H9c2 cell line were incubated with OA(5nM) and U0126(10μM) separately for 15-60min. p-ERK2^T183/Y185^(Green) was determined, Scale bar=20µm. (D), The transcriptional expression of mitochondrial dynamic related genes, after treatments as previously described in (C).

### 2.5 ERK2 is required for mitochondrial fragmentation in mouse hearts

The term mitochondrial dynamics has been used to represent a series of complementary fission and fusion events. The mammalian dynamin-related protein 1 (Drp1) and mitochondrial fission 1 (Fis1) proteins play a key role in fission[29, 30], while mitofusin 1 (Mfn1), mitofusin 2 (Mfn2) and opticatrophy 1 (OPA1) are required for fusion process[31, 32]. To understand how the nuclear accumulation of p-ERK2^T183/Y185^ associates with mitochondria dynamic regulation, we detected several genes responsible for mitochondrial dynamic regulation in H9c2 cells. As shown in Fig. 4D, U0126 pretreatment dramatically prevented the transcription of *Fis1*, albeit both fusion and fission genes are highly expressed after inhibitor treatments for 1h. Thus, this observation hinted that the nuclear translocation of p-ERK2^T183/Y185^ may promote the *Fis1* transcription to disturb the balance of mitochondrial fusion and fission.

As a key regulator of mitochondrial fission, integral outer membrane protein Fis1 recruits Drp1 onto the mitochondrial outer membrane. Completely assembled mitochondrial fission complex induces mitochondrial division. As p-ERK2^T183/Y185^ enhances *Fis1* expression, we next sought to address its effects on mitochondrial fragmentation. Mitochondrial fractions and cytosol suspensions were used for Western Bolting assay, respectively. As we expected, abundant Fis1 was detected in PP2Acα KO hearts, in both components. Concurrently, highly phosphorylated p-ERK2^T183/Y185^ and p-DRP1^S616^ were documented in their cytosolic fractions. Consequently, a significant increase of the Fis1 accumulation induced P62/SQSTM1, a mitophagy adaptor anchoring to mitochondria (Fig. 5A and 5B). In H9c2 cells, treatments of OA or U0126 coincided with the results observed in mouse hearts (Fig. 5C and 5D). Interestingly, the amount of mito-anchoring P62 expression increased along with an elevation of Fis1 level. These findings validated the fact that mitochondria fragmentation was highly related to p-ERK2^T183/Y185^ dependent Fis1 regulation.

**FIG. 5.**
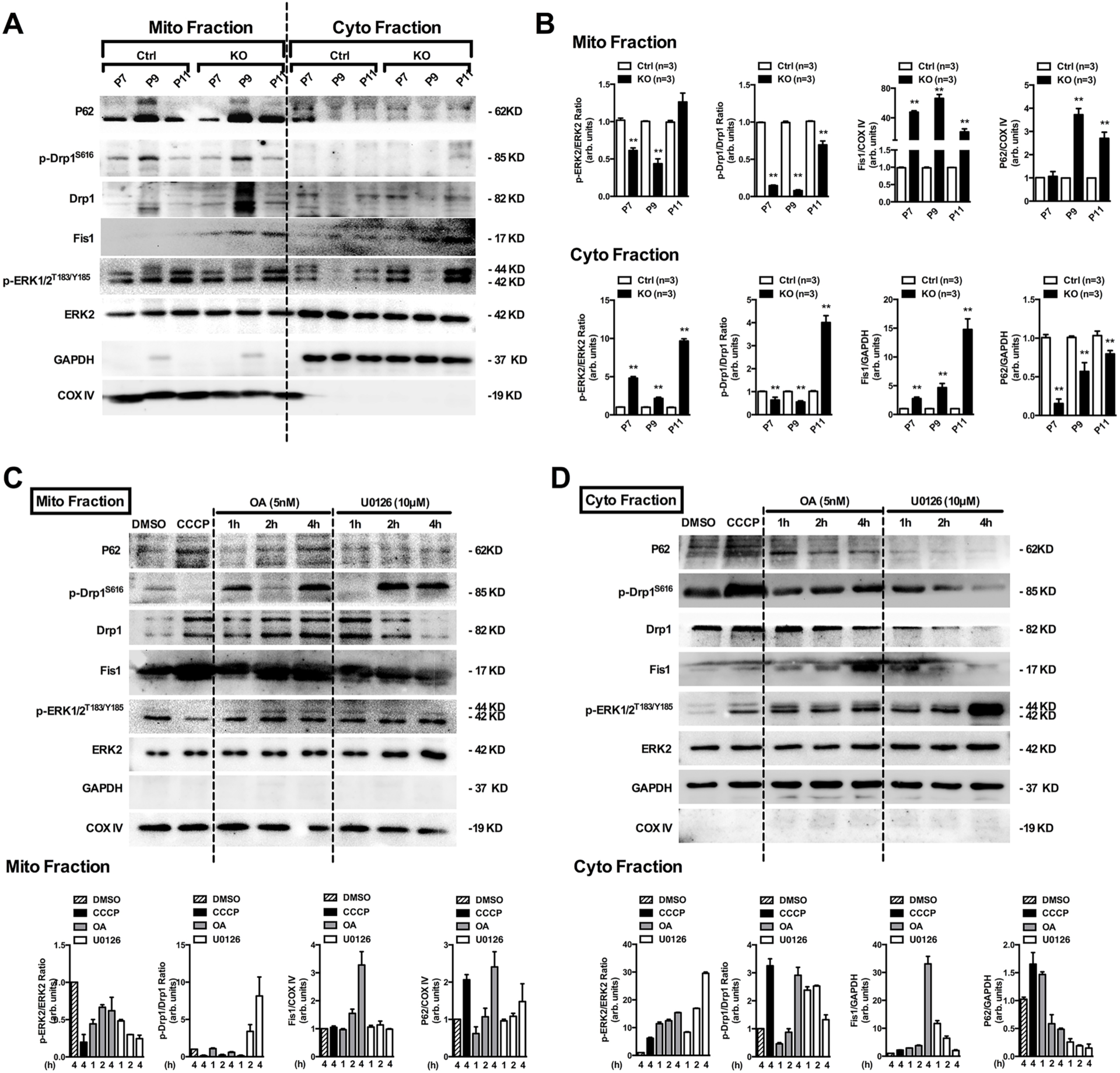
PP2Acα ablation/inhibition induced mitochondrial fission related mitophagy. (A), Isolated mitochondrial fraction from P7-P11 mice hearts and lysates were western blotted for indicated proteins. (B), Quantification from (A). (C), H9c2 cell were treated with OA(5nM) or U0126(10μM) for 4hr. Mitochondria were purified then subject to western blot using specific antibodies as in (A). (D), Quantification from (C). All data reported as mean±S.E., \**P*<0.05, \*\**P*<0.01 vs Ctrl. n=3 in each group.

### 2.6 Mitochondrial morphology and function could be manipulated via ERK2 phosphorylation regulation in H9c2 cells

Mitochondrial dynamics control mitochondrial function as well as their morphology. In our observations, disruption of mitochondrial membrane and dissipation of mitochondrial Δψm were not only evident in PP2Acα KO hearts (Fig. 1F) but also in H9c2 cells with incubation of either OA or U0126 for 1h to 4h, pointing to severe mitochondrial dysfunction in fusion-defective organelles (Fig. 6A). Abnormal respiratory function and increased ROS production in KO hearts further supported the dysfunction of mitochondria (Fig. 6B). In addition, CCK-8 analysis was used to evaluate the mitochondrial redox ability indirectly. Since the succinate dehydrogenase (SDH) in mitochondria is responsible for reducing exogenous cell counting kit-8 (CCK-8) reagents to purple formazan crystals[33]. The data demonstrated that the OA group presented increased mitochondrial viability (at 1h) followed by an alleviative decline. In other aspects, mitochondrial viability was continuously suppressed by U0126 inhibition. The viability of cells treated for 4h was reduced to 90% approximately (Fig. 6C).

**FIG. 6.**
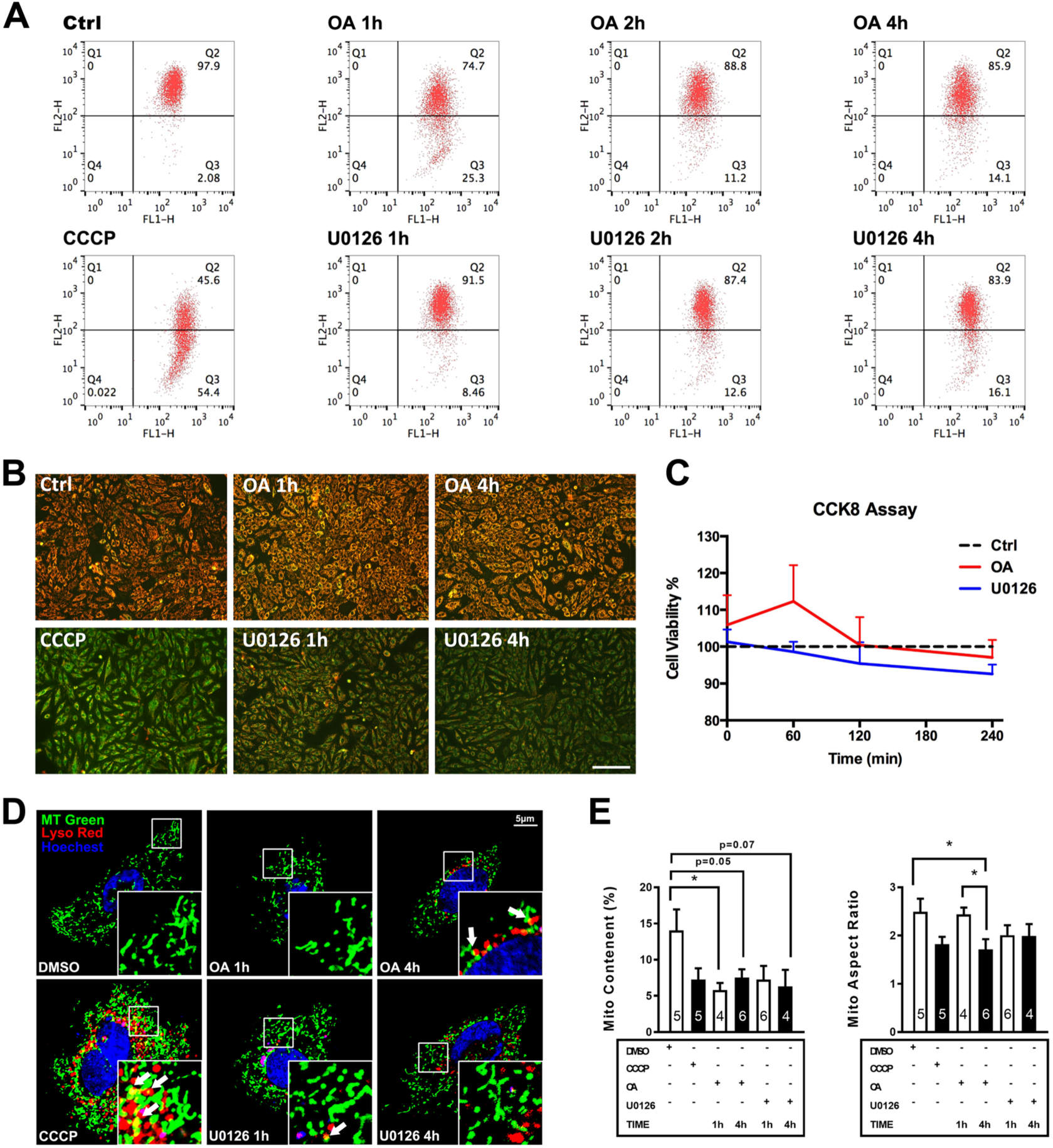
Mitochondrial structure and function were determined by the phosphorylation status of ERK2. (A), Flow cytometric measurements for MMP of isolated H9c2 mitochondria were labeled with JC-1. (B), Mitochondrial membrane potential (MMP) detected using fluor-microscopy. Scale bar=100µm. (C), Cell viability was analyzed using CCK8 assay. (D), Mitochondria of H9c2 co-stained with MitoTracker Green and LysoTracker Red. Wight arrow showed the sites interaction between mitochondria and lysosomes. (E), Quantitative mitochondrial continent (left) and mitochondrial aspect ratio (right). \**P*<0.05, \*\**P*<0.01 vs Ctrl. n= indicated number.

Elimination of impaired and superfluous mitochondria through mitophagy appears particularly important in terminally differentiated cardiomyocytes under physiological conditions and in response to pathological stresses. However, dysfunctional mitophagy might lead to cardiomyopathy and cell death. Thus, we investigated whether PP2A inhibition-induced the fusion-defect of mitochondria associates with dysfunction of mitophagy. We treated H9c2 cells with carbonyl cyanide m-chlorophenyl hydrazine (CCCP) to uncouple cell respiration within the mitochondria to trigger conventional Parkin-related mitophagy[34, 35]. The images of the H9c2 cells after 1h incubation with OA or U0126 showed obvious mitochondrial fragment and shortening (Fig. 7D, middle panel, top row). While a sustained OA treatment (4h) caused massive fragmentation and multiple conglomerates of mitochondria, as well as clumped lysosomes in the perinuclear region (colocalization indicated as white arrows shown in Fig. 7D, left panel, top row), indicating an enhancement of mitophagy. In comparison with OA treatment, the mito-lysosome interactions were rarely observed in persistent U0126-treated cells in spite of more and more lysosome aggregation in the nuclear periphery (Fig. 6D and 6E).

**FIG. 7.**
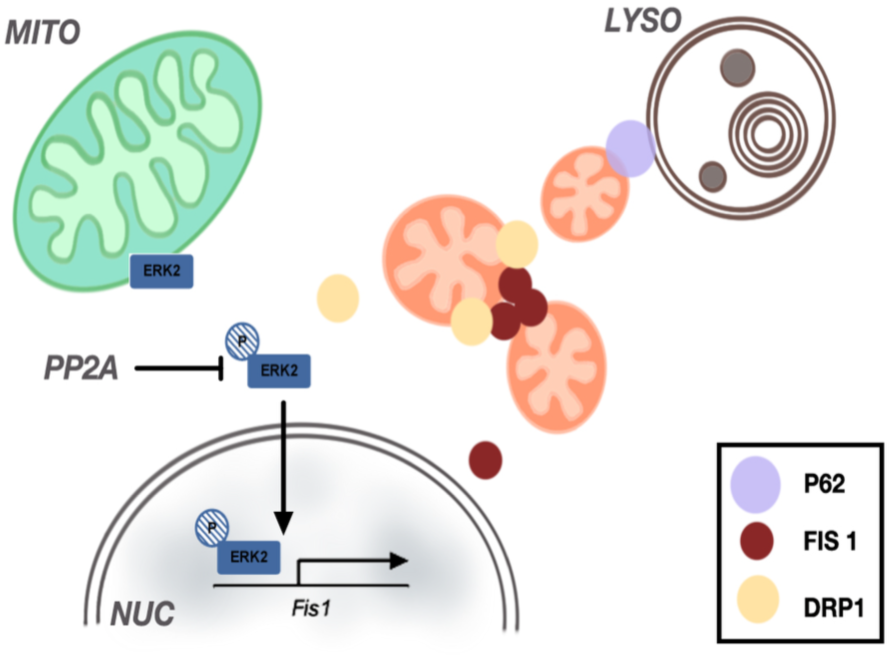
Schematic model of mitochondrial quality control by PP2Acα-dependent phosphorylation of ERK2. The schemata are represented in the summary of this study. Lacking PP2Acα caused a hyper-phosphorylation of ERK2, which translocate to nuclear and enhanced the expression of *Fis1*. Highly expressed Fis1 accumulated on the cardiac mitochondria, which recruited dynamic protein Drp1, accelerate the fission process. Extraordinary mitochondrial fragments trigger mitophagy consequently.

## 3 Discussion

In this study, we provide a new insight into the critical role of PP2A in the control of mitochondrial dynamics during postnatal heart development. A conditional PP2Acα abolishment induced hypertrophic cardiomyopathy, leading to early mortality in mice. Both *in vivo* and *in vitro* exams show a same response: PP2A deficiency causes intensified ERK2 phosphorylation. Subsequently over-phosphorylated ERK2 enriched into the nucleus, evoking *Fis1* expression. Exaggerated Fis1 recruits Drp1 to aggregate onto the outer mitochondrial membrane, thereby disturbing mitochondrial dynamics and shifting towards fission procedure. Eventually, fragmented mitochondria are degraded in the lysosome. Such severe damage of mitochondria and the resultant insufficient energy supplements might contribute to hypertrophic cardiomyopathy and the early mortality of PP2Acα KO mice.

In addition to the primary role of ATP generation, increasing evidence has shown that mitochondria are regarded as highly dynamic organelles. They are not only the powerhouse to create energy, but also play key roles in cell survival and death. It is well known that during the first few days of postnatal cardiac growth, the heart undergoes drastic changes in its metabolism and intracellular architecture as it switches from the placental circulation and anaerobic metabolism to the pulmonary circulation and oxidative phosphorylation. This postnatal transition is accompanied by proper coordination of mitochondrial fusion and fission. This transition process in the mouse heart with PP2Acα ablation seems to be impaired because of the disruption of mitochondrial dynamics, thus further contributing to the myocardial metabolic deficiency[36]. Therefore, our findings demonstrate that PP2A is closely associated with the maturation of intracellular energy pathways during postnatal heart development. Notably, mitochondrial fragmentation seems to be an early, upstream event in mitochondrial dysfunction that was documented in PP2Acα KO mice. A time-course profile of morphology for mouse hearts reveals that mitochondrial fragmentation occurred as early as P7 upon just one day after mutantial recombination was initiated. Concurrent results are evident in H9c2 cells by application of OA to inhibit PP2A.

Benefits from the application of iTRAQ-labeled phosphoproteomic and bioinformatics analysis, we identified the MAPKs as the predominantly altered signaling pathway besides several other components involved in regulation of the cell cycle or DNA damage and repair that are beyond in the scope of this study. Extensive studies have revealed that activation of MAPK subfamily is critically involved in a wide range of cardiac pathologies, such as cardiac injury, hypertrophy, heart failure and cardiac arrhythmias[6, 37, 38]. As known, the MAPKs are a group of protein Ser/Thr kinases consisting of three principal members, ERK1/2, JNK1/2/3, and p38 MAPKs. They are activated in response to a variety of extracellular stimuli and mediate signal transduction from the cell surface to the nucleus, thereby regulating various cellular processes including cell proliferation, differentiation, and survival. In the scenario of PP2Acα deletion, the phosphorylation of ERK2 respond rapidly with spatial and temporal features. Significant over-phosphorylation of ERK2 occurred at very early stage in PP2Acα KO as well as OA or U0126-treated H9c2 cells. One open question is why the phosphorylation status of ERK2 cannot last for a long period in our *in vitro* treatments. One possible explanation for the phosphorylation regression maybe a compensatory mechanism eliminated by autophosphorylated PP2A or other protein phosphatases. After U0126 application to inhibit MEK1/2, the phosphorylation level of ERK2 unanticipated rebound after 2h. This might be regarding adaptation or catabolism of U0126. Nevertheless, our findings demonstrate a potential pattern of how PP2A regulates MAPK signaling in time series.

Mitochondrial dynamics is the most important component for mitochondrial quality control, which is maintained and regulated by a number of key mitochondrial morphology proteins, including a series of dynamin motor-related GTPases to equilibrate fusion and fission. The present results indicate that PP2A deficiency-induced mitochondrial fragmentation is due to an imbalance favoring fission over fusion. For the fusion process, in PP2Acα KO hearts, we have observed an increased Mfn2 expression and mitochondria elongation (TEM images) at P7, but it has not been reproduced in H9c2 cells. There are no differences exhibited in Mfn1 expression. In other aspects, Fis1 is tightly correlated with the ERK2 phosphorylation in both animal and cultured cells. the fission process is promoted when overexpressing Fis1[39], while it is mild defected in Fis1^-/-^ cells[40]. Drp1 plays a key role in fission, which is mostly distributed in cytoplasm. Simply overexpressed Drp1 in cells have no significant effect on fission[41], but the regulation of Drp1 properties is more important. Drp1 recruitment to mitochondria increased prominently during events increased in fission[42]. More recently, some studies have reported an increased p-DRP1^Ser616^ induced by ERK2 and promote mitochondrial fission in the nervous system[43, 44]. In our PP2Acα KO mice, we have not observed increased phosphorylation modification on Drp1, but enhanced mitochondria translocation, altering fission significantly in mouse hearts.

There are multiple mechanistically distinct processes of dysfunctional mitochondria elimination that have been well described in numerous papers[45–47]. Recoverable mitochondria damage can be diluted via fusion-mediated reassembly. Irrecoverable damage will induce fission-mediated disassembly then removed by mitophagy. Mitophagy has recently been investigated and shown to be relevant to the cardiovascular system and as a novel therapeutic target for combating cardiovascular disease[36]. Hence, mitochondrial elongation at P7 in PP2Acα KO hearts was readily comprehended as a compensatory fusion. With increased mitochondrial impairment, consequently, massively segmented mitochondria were selectively engulfed by lysosomes. We revealed one facet of MAPK-mitophagy crosstalk the activation of oncogenic MAPK signaling promotes Fis1 expression and mitochondrial fission in mouse hearts. Additionally, mitochondrial dysfunction has been reported in MAPK-mutated children[48]. MAPK was required in mitophagy, which has been reported by several individual groups[43, 49, 50].

The mitophagy exhibited in PP2Acα KO hearts is similar to either CCCP induced Parkin dependent process or a mitochondrial depolarization related mitophagy as reported[51]. We also observed interaction between mitochondria and multivesicular bodies (MVBs) in PP2Acα KO hearts (Fig. S4), which were realized as a derived form of mitochondria-derived vesicles (MDV) and cargo to lysosome for degradation[52]. It is also a Parkin-dependent trafficking mechanism[53]. This implies that mitophagy in PP2Acα KO hearts is initiated by mitochondrial depolarization and is a Parkin-dependent pathway.

Inevitably, there are several regrettable flaws existed in this study. Firstly, it was difficult to provide a rescue experiment. We attempted to silence the MAPK signaling at the early stage via adenovirus intervention or administer the MEK agonist Curcumin with micropump implantation to attenuate the process of heart failure in PP2Acα KO mice. But we failed to achieve in such young mice due to technical obstacles. Next, the underlying mechanism by which ERK2 regulates *Fis1* has been characterized in this study, but no specific transcription factor is identified although predicted 3 putative binding sites on *Fis1* promoter are predicted. Our future studies will take on the indispensable task of finding a detailed mechanism.

Collectively, we provide a dynamic view of how PP2A induced post-translational modification controls mitochondrial dynamics and put a spotlight on MAPK signaling which is critically involved in the mitochondrial homeostasis in postnatal developing hearts.

## 4 Material and Methods

### 4.1 Animals

Animal experiments in this study were performed in accordance with the Guide for the Care and Use of Laboratory Animals as published by the US National Institutes of (NIH publication no.85-23, revised in 2011). All protocols were approved by the Laboratory Animal Care Committee at Nanjing Normal University (IACUC-1909002). All the mice used in this study were of the C57BL/6 background. Mice carrying PP2Acα alleles (*Ppp2ca^fl/fl^*) and α-MHC *Cre* transgenic mice were provided by Prof. Xiang Gao from the Model Animal Center of Nanjing University. All the animals were bred in a specific pathogen-free animal facility under standard conditions on a 12/12h light/dark cycle at a temperature of 22°C to 25°C and fed under *ad libitum*. Genotyping was performed by PCR analysis of genomic DNA that was extracted from mouse toes. Both males and females were used in the following experiments.

### 4.2 Transmission Electron Microscopy (TEM)

Freshly collected samples from the apex of the mouse left ventricle were dissected into 1-2 mm^3^ sections then immediately fixed with 2% glutaraldehyde, postfixed in 1% osmium tetroxide, and embedded in epoxy resin. Osmium tetroxide/uranyl acetate stained mouse heart thin sections (90 nm) were examined for mitochondrial ultrastructure by JEM-1400 (JEOL, Tokyo, Japan) at 1,500×∼30,000× direct magnifications. Several random fields were selected and all mitochondria within fields were calculated for amount, area and the aspect ratio using Image J.

### 4.3 Mitochondrial Flux Analyses

Mitochondria were isolated from the hearts of control and PP2Acα KO mice as previously described[54, 55]. Oxidative respiration rates (OCR) of the mitochondrial isolate containing 4 mg protein/well were measured using a Seahorse XF24 analyzer (Agilent Technologies, CA, USA) per the manufacturer’s protocol. Respiration was determined using pyruvate as a substrate. Following measurement of the basal respiration, maximal ADP-stimulated respiration was determined by exposing mitochondria to 4 mM ADP. Uncoupled respiration was evaluated following the addition of oligomycin (1 μM) to inhibit ATP synthase; this was followed by addition of the uncoupler FCCP (2 μM), and then the addition of antimycin A (0.2 μM).

### 4.4 High-Energy Phosphates Concentration

The dried samples were dissolved in 200 μl of mobile phase buffer (buffer A: 90 mM KH_2_PO_4_, 10 mM K_2_HPO_4_, 4 mM tetrabutylammonium sulfate/TBAS, pH 6.0). To remove insoluble material, the dissolved samples were centrifuged for 10 min at 20,000g. HPLC separation was carried out on a Shim-pack XR-ODS LC18 column (75 × 3.0mm, 5μm) using a Shimadzu HPLC system equipped with a SIL-20AC autosampler, CTO-20AC column oven, CBM-20A controller and UV-detector. 2μl of each sample was analyzed using a linear gradient of 100% buffer A to 60% buffer B (buffer B: 80 mM KH_2_PO_4_, 20 mM K_2_HPO_4_, 4 mM TBAS, 30% methanol, pH 7.2) over 30 min at 25°C. After each run, the column was eluted with 100% buffer B for 5 min and equilibrated again with buffer A for 5 min. The nucleotides were detected by their UV absorption at 260 nm.

### 4.5 Isolation of Mitochondria

Mouse hearts were collected, minced, and incubated with trypsin before homogenization with a glass/Teflon Potter Elvehjem homogenizer. Heart homogenates were centrifuged thrice at 800g for 10 min at 4°C and the supernatant was collected and again centrifuged thrice at 8,000g for 10 min at 4°C. Both the pellet and the supernatant were collected. The pellet was again washed and centrifuged thrice at 8,000 g for 10 min at 4°C to obtain normal-size mitochondria, which were resuspended for analyses. The 8,000 g supernatant was centrifuged thrice at 16,000 g for 10 min at 4°C to pellet smaller fragmented mitochondria.

### 4.6 Flow Cytometric Analyses of Isolated Mitochondria

Mitochondrial membrane potential (Δψm) was assessed by JC-1 dye (Beyotime Biotechnology, Shanghai, China), by flow cytometry, and the data were collected using a BD-FACS (Becton Dickinson, NJ, USA). Analyzed by FlowJo (v10.0.7 for Mac, Treestar, CA, USA).

### 4.7 Phosphoproteomics

*Sample Preparation:* Tissue samples were prepared using Ready Prep Protein Extraction kit (Bio-RAD, CA, USA). Approximately 400 mg of protein/sample was used for quantitative phosphoproteomic profiling. In order to ensure adequate coverage of phosphosites[56], 3 biological replicates (a total of 6 samples) including 3 Ctrl and 3 KO samples were collected from 3 independent cultures.

*Proteolytic Digestion and Peptide Labeling by iTRAQ Reagents:* Each protein sample was reduced with 10 mM DTT for 1 h, followed by alkylation with 40 mM iodoacetamide for 1 h under dark conditions. Samples were diluted with 50 mM ammonium bicarbonate to a less than 1 M final urea concentration before protein digestion with trypsin (Promega, WI, USA) at a mass ratio of 1:20 (trypsin: protein) overnight at 37°C Following tryptic digestion, peptide samples were desalted using MonoTip C18 (Shimadzu Biotech, Kyoto, Japan). The eluted peptides were dried in a SpeedVac and then labeled with iTRAQ reagents according to the manufacturer’s instructions. Each of the six samples was labeled separately with 4 vials of iTRAQ isobaric reagent (114, 115, 116, 117, 119 and 121), respectively. After incubation for 2 h at room temperature, the reaction was stopped by acidification with formic acid (1%). 6 iTRAQ-labeled peptide samples were then combined and desalted using MonoTip C18 (Shimadzu Biotech, Kyoto, Japan). The eluted peptides were dried in a SpeedVac.

*Enrichment of Phosphopeptides and LC-MS/MS Analysis:* Phosphopeptide enrichment was performed using Titansphere Phos-TiO kit (Shimadzu Biotech, Kyoto, Japan) according to the manufacturer’s instructions. Elution of phosphopeptides was carried out using 50mL of elution buffer each, three times. Combined eluents were acidified with 100mL of 2.5% trifluoroacetic acid. The samples were then desalted with the MonoTip C18 (Shimadzu Biotech, Kyoto, Japan) and resuspended in 0.1% formic acid prior to analysis by LC-MS/MS. Samples were analyzed on a Prominence nanoflow LC system (Shimadzu Biotech, Kyoto, Japan) connected to a LCMS-IT-TOF mass spectrometer (Shimadzu Biotech, Kyoto, Japan)[57]. iTRAQ labeled ions were fragmented in IT (IonTrap). CID fragmentation ions were detected in the TOF. MS2 spectra were searched with Mascot engine using the following criteria: database, Swiss-Prot, mouse; enzyme, trypsin; miscleavages, 2; static modifications, carbamidomethylation of cysteine (+57.021 Da), iTRAQ8-plex modification of peptide N-terminal (+304.205 Da), iTRAQ8-plex modification of lysine (+304.205 Da); variable modifications, oxidation of methionine (+15.995 Da); phosphorylation of serine, threonine and tyrosine (+79.966 Da); iTRAQ8-plex modification of tyrosine (+304.205 Da); precursor ion tolerance, 25 ppm; fragment ion tolerance, 0.05 Da. The dataset was filtered to give a <1% false discovery rate at peptide level using the target decoy method. The phosphorylation sites were determined using PTM Finder Software (Shimadzu Biotech, Kyoto, Japan).

*iTRAQ Quantification:* Abundance ratios (KO/Ctrl) were quantified by Proteome Discoverer Software (Thermo-Fisher Scientific, CA, USA) via the quantification of iTRAQ labeled peptides[58]. To minimize contaminating near isobaric ions, only the peptides with isolation specificity more than 75% were quantified. For redundant peptides, the KO/Ctrl ratio was calculated from the pair with the highest summed reporter ion intensity. The phosphopeptide ratios were used as the basis for the calculation of the mean value for each peptide across all three biological replicates. An unpaired *t*-test was used to determine whether changes in phosphopeptide abundances were significant.

### 4.8 Co-immunoprecipitation

For immunoprecipitation experiments, mice samples were flash-frozen in liquid nitrogen and ground into a fine powder. Samples were resuspended in homogenization buffer (0.32 M sucrose, 2.5mM EGTA, 5mM EDTA, 50mM Tris, and 10mM NaCl, pH 7.4) and further homogenized by mechanical agitation with a Dounce homogenizer. NP-40 (1% final) or RIPA (0.1% SDS, 1% Triton X-100) was added to each sample, and the samples were sonicated. Lysates were centrifuged at 3000g for 15 min.

The cells were lysed in PBS (pH 7.4). Lysates were incubated with the indicated antibody (1:50, overnight, 4 °C), and the protein-antibody complex was precipitated by centrifugation after incubation with protein anti-IgG2b agarose beads (2hr, 4 °C). The immunoprecipitated material was washed twice in PBS and resuspended in Native loading buffer (BN2003; Invitrogen, CA, USA), and loaded on 4%–12% PAGE gels.

### 4.9 Isolation of Cardiomyocytes

Ventricular myocytes were isolated from newborn mice at P7, P9, and P11 as previously described[59]. Briefly, after anesthesia with 0.1ml sodium pentobarbital (i.p. 40mg.kg^-1^), the heart was rapidly excised, cannulated and then mounted on a Langendorff apparatus. Following a brief period of perfusion with oxygenated calcium-free Tyrode’s solution (140 mM NaCl, 5.4 mM KCl, 1 mM MgCl_2_, 10 mM HEPES, 10 mM Glucose, pH 7.4), the heart was perfused with oxygenated Tyrode’s solution containing 0.3 mg/ml collagenase B (Sigma-Aldrich, MO, USA ) and 0.6% BSA at 37°C for 3 min, and then switch to high K^+^ (HK) solution (120 mM L-Glutamic acid, 80 mM KOH, 20 mM KCl, 1 mM MgCl_2_, 0.3 mM EGTA, 10 mM HEPES, 10 mM Glucose, pH 7.4) for 5 min. In succession, the ventricles were minced and gently pipetted by Pasteur pipette. Isolated cells were filtered through nylon mesh (200 µm) and then resuspend in HK solution incubated at 37°C for 1 hour.

### 4.10 Cell Culture

H9c2 cells (American Type Cell Cutler, Manassas, VA, USA) were maintained in Dulbecco’s modified Eagle’s medium (DMEM) supplemented with 10% fetal bovine serum, 2 mM L-glutamine, 100 U/ml penicillin, and 100 mg/ml streptomycin. Cells were seeded in 12- and 6-well cell culture plates at 37°C under a 5% CO2 humidified environment for RNA and protein assays, respectively, or in 96-well cell culture plates for the CCK8 assay. In all experiments, the cells were treated for the indicated time intervals in serum-free media with various concentrations of Okadaic Acid (OA) and inhibitor U0126. Stock solutions of OA were prepared fresh in dimethyl sulfoxide (DMSO) just prior to each experiment, where the DMSO concentration in all the treatment did not exceed 0.05 % (v/v).

### 4.11 Confocal Microscopy

Live-cell imaging used a Nikon Ti confocal microscope equipped with a 60× or 100× 1.3NA oil immersion objective lens. For visualization of mitochondria and measurement of mitochondrial membrane potential, cells were stained with 200 nM MitoTracker Green, 200 nM of tetramethylrhodamine, ethyl ester (TMRE), and 10 mg/ml Hoechst at 37°C for 30 min. For detection of parkin aggregation, cells were counter stained with MitoTracker Green and Hoechst. For assessment of lysosomal engulfed mitochondria, cells were stained with 50 nM LysoTracker Red, MitoTracker Green, and Hoechst. Images were analyzed by Image J.

### 4.12 Image Analysis

Mitochondrial aspect ratio (the ratio of length/width) and content (% of mitochondrial area compared to whole-cell area) were quantified using Image J. Mitochondrial depolarization was calculated as % of the area of green mitochondria compared to that of all the mitochondria visualized on MitoTracker Green and TMRE merged images; data are presented as green/(green + yellow mitochondria)× 100%. Parkin aggregation was calculated as % of cells with clumping mCherry-Parkin compared to all the cells. Lysosomal engulfment of mitochondria was calculated by counting the number of colocalized lysosomes and mitochondria per cell detected by confocal colocalization of LysoTracker Red and MitoTracker Green.

### 4.13 Transfection

Specific rat PP2Acα silencer designed interfering small RNAs (siRNA) and scrambled control siRNA were purchased from Bioworld Technology, Inc (Nanjing, China). The effectiveness of three candidate siRNAs was confirmed by western blot. H9c2 cells were cultured in 6-well plates, si-PP2A Cα-3 was transfected in cells by Lipofectamine 3000 (Invitrogen, CA, USA). Cells were collected at 48, 60, and 72 hours after transfection.

### 4.14 RNA Expression Analysis

Total RNA was extracted from the left ventricle using Trizol reagent (TaKaRa, Shiga, Japan). cDNA was synthesized using the PrimeScript™ 1st Strand cDNA Synthesis Kit (TaKaRa, Japan) according to the manufacturer’s protocol. Relative mRNA expression levels of genes of interest were quantified by RT-PCR using the SYBR® Premix Ex Taq™ kit (TaKaRa, Shiga, Japan) and performed in StepOnePlus™ Real-time PCR Detection System (Applied Biosystems, CA, USA).

Primers used as follows:

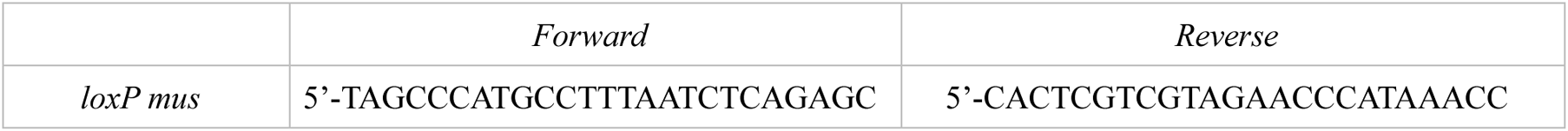
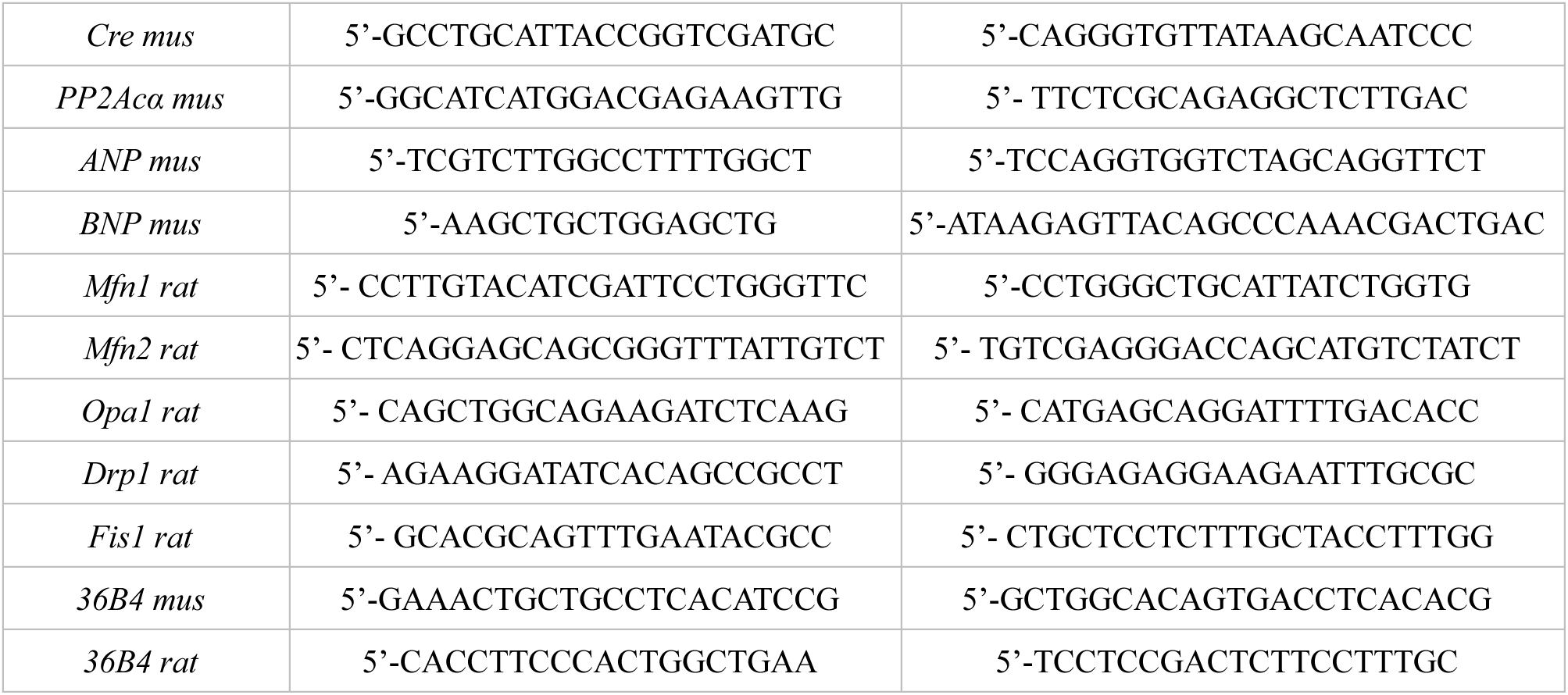

### 4.15 Antibodies

For Western blotting, the primary antibody against PP2Acα (610556) was from Becton Dickinson Company, and primary antibodies against PP2Ac (1:2000, #2259), Drp1 (1:1,000, #14647), p-Drp1(1:1000, #4494), ERK1/2 (1:1000, #4695), p-ERK1/2 (1:1000, #4370), JNK(1:1000, #9252), p-JNK(1:1000, #4668), P38(1:2000, #8690), p-P38(1:2000, #4511), GAPDH(1:2000, #5174) COX(1:2000, #4850)were from Cell Signaling Technology. The primary antibody against Fis1 (1:500, sc-84687) was from Santa Cruz Biotechnology. Horseradish peroxidase (HRP) linked secondary antibodies anti-mouse IgG (1:3000, #7076) and anti-rabbit IgG (1:3000, #7074) were from Cell Signaling Technology.

Immunoprecipitation antibody IgG2b (1:100, sc-537591) was from Santa Cruz Biotechnology. For immunohistochemistry, primary antibodies against α-actinin (1:200, ab9465) were from Abcam. Alexa Fluor-conjugated secondary antibodies anti-mouse IgG (1:400, A-11029) and anti-rabbit IgG (1:400, A-11035) were from Invitrogen.

### 4.16 Statistical Analysis

Grouped data were expressed as mean ± SEM. The Student’s *t*-test was performed for analyses of the data using GraphPad Prism (GraphPad, CA, USA) for Mac OS X. Statistical significance was set at \**P*<0.05 (compared with control).

## Acknowledgments

This project was supported by the National Science Foundation of China (Nos. 30570662, 30871228 and 31171302) and by the grant of Priority Academic Program Development of Jiangsu Higher Education Institutions (164320H106), the Major Program of Educational Commission of Jiangsu Province (16KJA180001). The PP2Acα and α-MHC-Cr*e* transgenic mice were provided by Prof. Xiang Gao (Nanjing University, China)

## Author contribution

D.D. and Z.Z. designed experiments. D.D., Y.Z., L.L., H.F., and T.J. performed experiments and analyzed data with supervision by Z.Z.. D.D. wrote the manuscript. D.D. and Z.Z. edited the manuscript. All authors commented on the manuscript. Z.Z. supervised all aspects of the work.

---

**FIG. S1.**
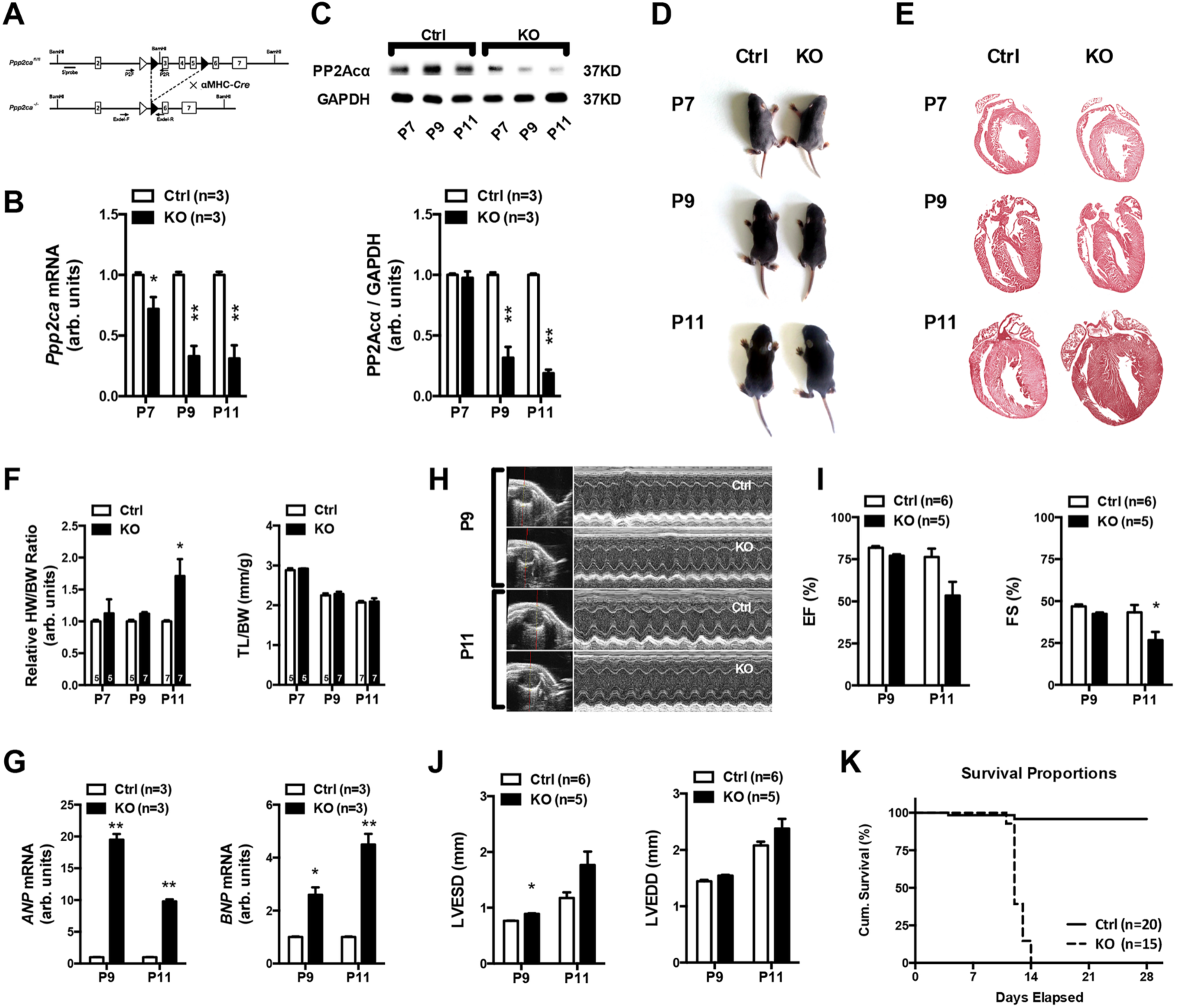
Cardiac-specific PP2Acα deletion leads to hypertrophic cardiomyopathy. (A) Schematic view of the targeted *Ppp2ca* knockout strategy. Exons are represented by open boxes in which the exon number is indicated. Black triangles denote loxP sites. *Ppp2ca*^-/-^ mice were generated by crossing *Ppp2ca ^fl/fl^* mice to cardiac-specific transgenic aMHC-*Cre* mice. (B), Cardiac total RNA was analyzed for *Ppp2ca.* Expression was normalized to *GAPDH.* (C), Tissue lysates from hearts in (B) were analyzed for PP2Acα expression. (D), Gross morphology of Ctrl and PP2Acα KO mice from P7 to P11. (E), H&E staining of a sagittal section of Ctrl and PP2Acα mutant hearts from P7-P11. (F), Hypertrophy in PP2Acα KO mice was indicated by increased ratios of heart weight to body weight (HW/BW). (G), The hypertrophic markers highly expressed from P9. (H-J), Cardiac hemodynamic. M-mode echocardiography obtained from the parasternal short-axis view at the mid-papillary level, in Ctrl and PP2Acα KO mice, at P9 and P11 (H). PP2Acα deletion impaired the contraction function of hearts (I and J). (K), The survival Kaplan-Meier curve describes the lifespan of Ctrl and PP2Acα KO mice, *P*<0.01. All data reported as mean±S.E., \**P*<0.05, \*\**P*<0.01 vs Ctrl group. n= indicated numbers of mice.

**FIG. S2.**
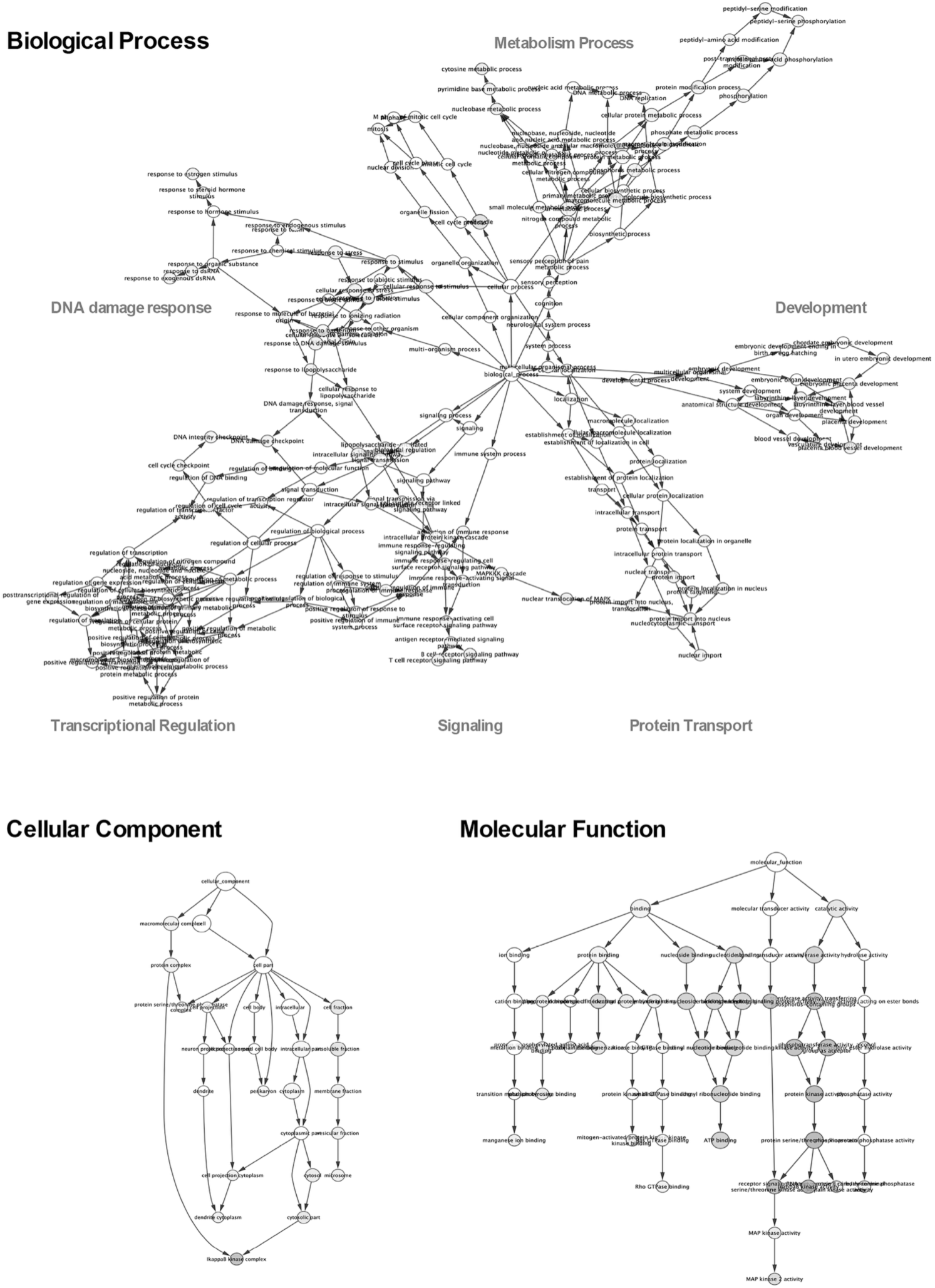
GO annotations of phosphorylated proteins identified by proteomics. Enrichment of identified phosphorylated protein complexes for biological process, cellular component, and molecular functional terms.

**FIG. S3.**
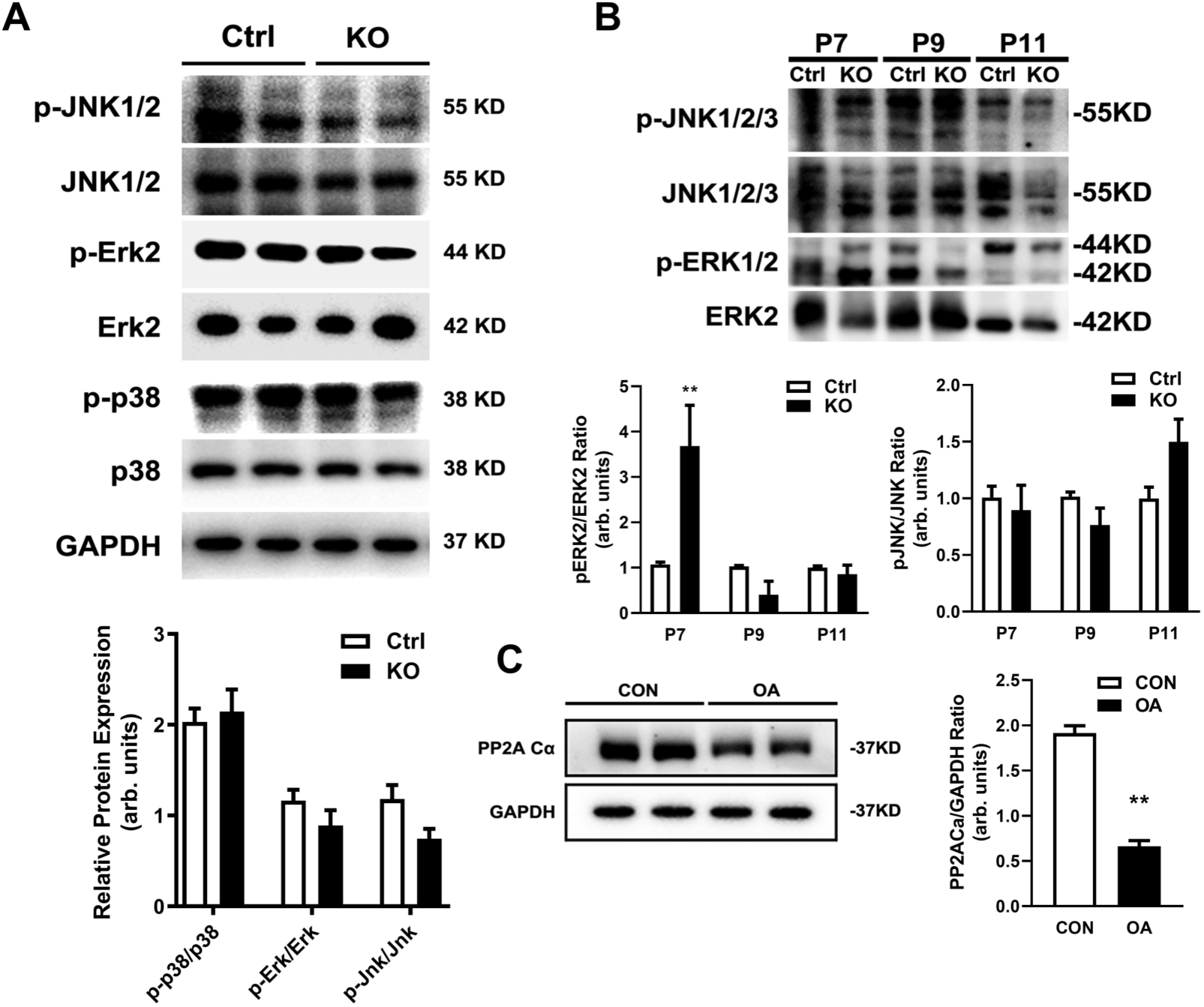
Effect of PP2Acα deficiency on three MAPK members. (A), expression and phosphorylation level of three MAPK members at postnatal day11. (B), The performance of phosphorylation response of MAPK members on a time scale. (C), Okadaic Acid inhibits PP2A activity without any impact on PP2A content.

**FIG. S4.**
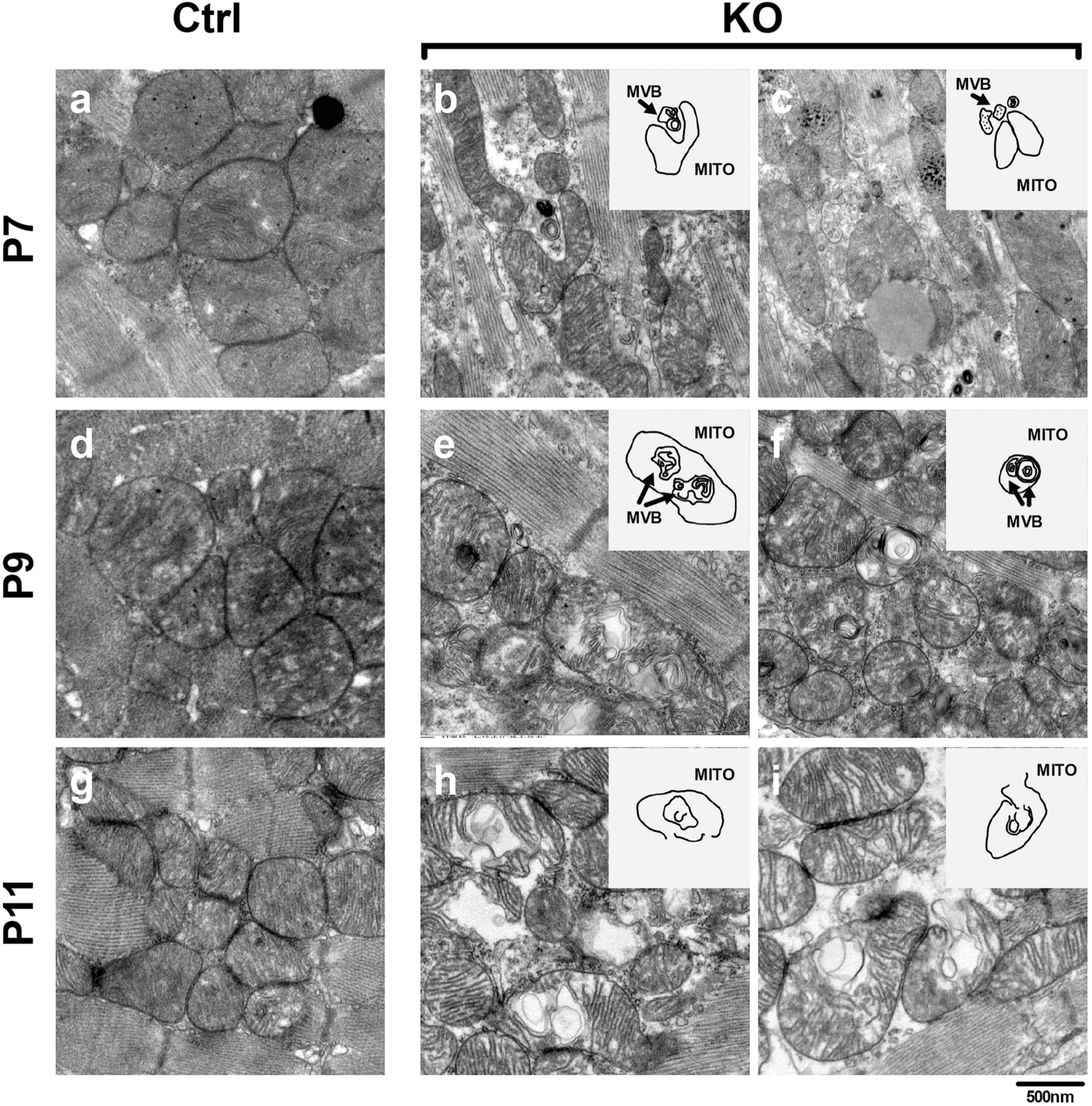
Electron micrographs indicated that multivesicular bodies (MVBs) participated in mitochondria-derived mitophagy in PP2Acα deficient hearts. At postnatal day7, autophagic vacuoles displaying an advanced degradation of multilamellar lysosomal bodies in KO heart (b,c), At postnatal day P9, mitochondria of PP2Acα KO heart appeared rounded and tightly contact with MVBs (e,f) and at the end stage, both MVB nano vesicular and mitochondria are collapsed in KO mice (h,i) Scale bar:500nm

